# *Aspergillus fumigatus* drives tissue damage via iterative assaults upon mucosal integrity and immune homeostasis

**DOI:** 10.1101/2021.11.09.468003

**Authors:** Uju Joy Okaa, Margherita Bertuzzi, Rachael Fortune-Grant, Darren D. Thomson, David L. Moyes, Julian R. Naglik, Elaine Bignell

## Abstract

The human lung is constantly exposed to *Aspergillus fumigatus* spores, the most prevalent worldwide cause of fungal respiratory disease. Pulmonary tissue damage is a unifying feature of *Aspergillus*-related diseases; however, the mechanistic basis of damage is not understood. In the lungs of susceptible hosts *A. fumigatus* undergoes an obligatory morphological switch involving spore germination and hyphal growth. We modelled *A. fumigatus* infection in cultured A549 human pneumocytes, capturing phosphoactivation status of five host signalling pathways, nuclear translocation & DNA binding of eight host transcription factors, and expression of nine host response proteins over six time points encompassing exposures to live fungus and the secretome thereof. The resulting dataset, comprised of more than 1000 data points, reveals that pneumocytes mount differential responses to *A. fumigatus* spores, hyphae and soluble secreted products via the NF-kB, JNK, and JNK + p38 pathways respectively. Importantly, via selective degradation of host pro-inflammatory (IL-6 and IL-8) cytokines and growth factors (FGF-2), fungal secreted products reorchestrate the host response to fungal challenge as well as driving multiparametric epithelial damage, culminating in cytolysis. Dysregulation of NF-kB signalling, involving iterative stimulation of canonical and non-canonical signalling, was identified as a significant feature of host damage both *in vitro* and in a mouse model of invasive aspergillosis. Our data demonstrate that composite tissue damage results from iterative exposures to different fungal morphotypes and secreted products and suggest that modulation of host responses to fungal challenge might represent a unified strategy for therapeutic control of pathologically distinct types of *Aspergillus*-related disease.

**IMPORTANCE:** Pulmonary aspergillosis is a spectrum of diseases caused primarily by *Aspergillus fumigatus*. This fungus is ubiquitous in the environment and grows as a mold, which harbors and disperses spores into the environment. Like other airborne pathogens, the lung mucosa is the first point of contact with the fungus post inhalation. The outcome and severity of disease depends on the host-fungal interaction at the lung interface. We studied how the human lung interacts with spore, germ tube and hyphae growth forms to understand the sequence and dynamics of the early events, which are critical drivers of disease development and progression. Our work is significant in identifying, in response to fungal secreted products, non-canonical NF-kB activation via RelB as being a driving factor in fungus-mediated lung damage. This process could be modulated therapeutically to protect the integrity of infected lung mucosae.

## Introduction

*Aspergillus fumigatus* is an opportunistic fungal pathogen and a major cause of human lung disease. *A. fumigatus* spores are common amongst the airborne microflora and are associated annually with ∼200,000 human fatalities, occurring in severely immunocompromised patients including organ, or stem cell transplantees, HIV/AIDS or underlying chronic pulmonary disease (1). In asthma, bronchiectasis and cystic fibrosis settings inhalation of *A. fumigatus* spores also causes millions of debilitating respiratory malfunctions including allergic bronchopulmonary aspergillosis (ABPA) and chronic pulmonary aspergillosis (CPA) (1). With the emergence of new risk factors such as severe acute respiratory syndrome coronavirus 2(SARS-CoV-2), chimeric antigen receptor T-Cell (CAR-T cell) therapy, intensive care unit (ICU) stay, Aspergillus related morbidity and mortality remains increasingly high (2).

During contact with the pathogen, airway epithelial cells (AECs) are iteratively exposed to *A. fumigatus* spores, swollen spores, elongated hyphae and secreted fungal products, all of which elicit host responses that can culminate in diverse disease pathologies (3). In addition to physical invasion of lung tissues, *A. fumigatus* pathogenicity is also driven by secreted factors produced by the fungus, such as proteases and immunotoxins, which become expressed downstream of an obligatory morphogenetic switch from a spore to a hyphal form (4-9). The iterative interactions of *A. fumigatus* spore and hyphal forms with host cells is an integral feature of the host-pathogen interaction; however, our current understanding of these interactions has been compiled from disparate, mostly single time point, studies which have used different types of AECs and varying *A. fumigatus* morphotypes (8). Such approaches overlook the effects of iterative host exposure to multiple fungal morphologies that occurs during the sustained host-pathogen interaction during epithelial colonisation. Amongst other fungal pathogens that colonise and infect host mucosa, evidence of morphotype-specific activation of host responses has been demonstrated in response to *Candida albicans* challenge (4, 5). Hypothesising that sequential, morphotype-specific host responses to *A. fumigatus* challenge drives composite damage during *A. fumigatus* infection, we sought to capture the dynamic epithelial responses to *A. fumigatus* as a means to study the relative contributions of pathogen and host activities to epithelial damage.

## Materials and methods

### Culture of airway epithelial cells (AECs)

The A549 human alveolar type II-like carcinoma cell line (American type culture collection, ATCC ®CCL-185™) (10) was cultured in Roswell Park Memorial Institute 1640 Medium (RPMI 1640) plus glutamine, 10% fetal bovine serum (FBS) and 1% penicillin-streptomycin (10,000 units penicillin and 10 mg streptomycin/mL) at 37°C and 5% CO_2_ in a humidified atmosphere. Serum was omitted from the culture medium 18-24 h prior to fungal challenge and fungal challenges were carried out in serum-free RPMI 1640 plus glutamine and 1% penicillin-streptomycin. For transwell experiments, Calu-3 bronchial adenocarcinoma cells (ATCC ® HTB-55™) (11) were cultured in DMEM-F12 (1X) (Thermo-Fisher Scientific) supplemented with 10% FBS and 1% penicillin-streptomycin. For chemical inhibition of signalling pathways, 1 h prior to fungal challenge, confluent epithelial cell monolayers were treated with the following inhibitors at concentrations previously determined (5): SP600125 (Primary target: JNK1, JNK2, JNK3; 10 μM, Calbiochem), SB203580 (Primary target: p38; 10 μM, Calbiochem) and BAY11-7082 (Primary target: IκB-α; 2 μM Calbiochem).

### Fungal strains and culture

*A. fumigatus* isolates used in this study are listed in Supplementary Data 1. *A. fumigatus* was grown at 37°C for 3-4 days on *Aspergillus* complete medium (ACM) agar (12). Spores were harvested using sterile water, enumerated and resuspended at the desired concentration in serum-free RPMI 1640 medium. Unless otherwise stated, in *in vitro* infection assays, a final concentration of spores of 1×10^5^ spores/ml was used for detachment and transwell experiments, while 1×10^6^ and 1×10^7^ spores/ml were used for cytotoxicity and host signalling respectively. Heat-killing was achieved by incubation for 40 min at 90°C. To generate specific morphotypes, spores were pre-incubated in serum-free RPMI 1640 medium at 37°C, for 4 h to generate swollen conidia, and for 8 or 12 h respectively to generate young germlings or mature hyphae. To make culture filtrates (CF), freshly harvested conidia were resuspended at a cell density of 1×10^6^ spores/ml in RPMI 1640 medium, and incubated with shaking at 200 rpm and 37°C for 16 (CF^16^) or 48 h (CF^48^). The culture suspension was filtered through sterile Miracloth (Calbiochem) and a 0.22 µm filter, then stored at -20°C until use. Unless otherwise stated, challenges of the epithelial monolayers were carried out using a 5 fold dilution of the CF^48^ in serum-free RPMI 1640 medium.

### Microscopy

#### Microscopical analysis of challenged monolayers

Upon removal of culture media and a triple wash with PBS, *A. fumigatus* challenged monolayers were incubated with 1 ml of serum-free RPMI 1640 medium containing 1 µg/ml Concanavalin A conjugated to FITC (Con-A, Sigma) for 30 min at 37°C and 5% CO_2_. Widefield epifluorescence microscopy was performed using a Nikon Eclipse TE2000-E microscope (Nikon Instruments, Europe BV, UK) and a Nikon PlanFluor 20x/0.75 NA objective lens. A CoolLED PreciseExcite system (CoolLED, Andover, UK) was used with 470 nm and 550 nm LED arrays for FITC and tdTomato excitation, respectively. Images were captured with an ORCA-ER CCD camera (Hamamatsu, Welwyn Garden City, UK) driven by MetaMorph software v7.7.6.0 (Molecular Devices, Sunnyvale, CA, USA).

#### Comparison of fungal biomass

Fungal biomass of 10^4^ spores/ml of selected *A. fumigatus* isolates was assessed by microscopy at 4, 8, 12, 16 and 20 h of growth in supplemented RPMI-1640. Cumulative fungal length after 24 h of growth in supplemented RPMI-1640 exceeded the field of view boundaries and therefore was not measurable at the time. Widefield epifluorescence microscopy of *A*.*fumigatus* was performed using a Nikon Ti-S microscope (Nikon Instruments, Europe BV, UK) and a Nikon PlanFluor 20x/0.75 NA objective lens. A CoolLED PreciseExcite system (CoolLED, Andover, UK) was used with a 550 nm LED array for tdTomato excitation. Images were captured with Images were captured with a a Brightline LED-DA/FI/TR/CY5-A-000-ZERO multiband filter and an ORCA-ER CCD camera (Hamamatsu, Welwyn Garden City, UK) driven by Nikon Elements (v.5.11.01). Cumulative filament length (including branches) of 7-10 hyphae for each strain and time-point was measured in FIJI using the line summation function (13).

### Analysis of A549 monolayer integrity by detachment

Monolayers were challenged with 10^5^ *A. fumigatus* spores or CF. Following co-incubation, monolayers were washed 3 times with PBS, fixed with 4% formaldehyde in PBS, and permeabilized with 0.2% Triton-X100. Nuclei of adherent A549 cells were stained with 300 nM DAPI. DAPI fluorescence in adherent epithelial cells was excited with a CoolLED PreciseExcite 380 nm LED array in combination with a Nikon UV-2A filter cube, which collected the DAPI emission. Images were acquired as stated above, where at least three fields of view, per experimental well were taken. Images were then processed and quantified using an in-house “DAPI Counter” macro written for FIJI (Supplementary Data 2). Experiments were performed in biological triplicates with 3-5 technical replicates, whereby a technical replicate is a different field of view for the enumeration by microscopy. Each datapoint on figures represents a technical replicate of each of the biological replicates.

### Epithelial cytotoxicity

Epithelial cytotoxicity was determined by quantification of lactate dehydrogenase activity in A549 culture supernatants at 24 h post fungal challenge using the Cytox 96 Non-Radioactive Cytotoxicity Assay kit (Promega) according to the manufacturer’s protocol and using recombinant porcine LDH enzyme (Sigma-Aldrich) for derivation of a standard curve. Experiments were performed in biological triplicates with 1-4 technical replicates, whereby a technical replicate is a different infection well. Each technical replicate was measured for LDH twice and each data point on figures represent the average of the LDH measurements for each techical replicate of each of the biological replicates.

### Measurement of Trans-epithelial electrical resistance (TEER)

Calu-3 cells were seeded in DMEM-F12 (1X) in trans-well inserts (Scientific Laboratory Supply (SLS) placed in a 12 well tissue culture plates (Scientific Laboratory Supply) containing 2 mL of supplemented DMEM-F12. Media was changed every 2 days and TEER was measured every 4 days using a World Precision Instrument Evom2 epithelial voltohmeter and STX-2 electrode until it reached at least 1000 Ωcm^-2^ (∼11 to 13 days). TEER measurements were taken prior to infection with live spores or CF^48^ (TEER 0 h) and following 24 h of incubation (TEER 24 h). The decrease of TEER upon fungal challenge was calculated as the difference between TEER 24 h and TEER 0 h, normalised to fungal viable counts and expressed as fold reduction in TEER relative to PBS challenge. Experiments were performed in biological triplicates with 2 technical replicates, whereby a technical replicate is a different infection well. Each datapoint on figures rapresents a technical replicate for the biological replicates.

### Quantification of cytokine expression

To quantify cytokine secretion, A549 cell culture supernatants were collected at 24 h post infection with *A. fumigatus* spores or CF. Two distinct experimental approaches were then taken, the first adopting an unbiased semi-quantitative approach based on the HXL cytokine profile kit from R&D systems (Bio-Techne). Cell culture supernatants were diluted 1:3 and procedure was carried out according to the manufacturer’s instructions, aquiring data using a ChemiDoc MP imaging system (BIO-RAD). The relative amount of each cytokine was calculated by deriving the mean pixel density using ImageJ and micro-array profile plugins (Bio-Techne) and normalised relative to the uninfected control. To normalise between biological replicates the data were ranked according to protein quantity and abundant proteins defined as those represented in the top or bottom quantiles were considered as being significantly up- or downregulated. In a second experimental approach, a total of 25 μl of the cell culture supernatant was used to determine cytokine levels using the luminex high performance cytokine assay (Bio-Techne, United Kingdom) according to the manufacturer’s specifications. AECs stimulated with 50 ng/ml TNF-α served as a positive control of cytokine induction. Data was analyzed using Logistic 5PL analysis within the Bio-Plex Manager 6.1 software suite (Bio-Rad, United Kingdom).

### Quantification of phosphorylated signalling molecules in AECs

Challenged A549 cells were lysed with modified RIPA buffer (Supplementary Data 3) and protein concentration was determined using a bicinchoninic acid (BCA) protein assay (Thermo Fisher Scientific). Phospho-p38 (p-p38), phospho-Jun N-terminal protein kinase (p-JNK), phospho-extracellular signal-regulated kinases 1 and 2 (pERK1/2), phospho-IκBα (p-IκBα), and phospho-Akt (p-Akt) were quantified in 10 μg of total protein using the Bio-Plex Pro magnetic phosphoprotein detection assay (Bio-Rad, United Kingdom). Phosphoprotein expression was normalized to β-actin and expressed as fold-change relative to the uninfected (UI) control.

### Quantitation of transcription factor binding

Nuclear proteins were isolated from challenged A549 cells using a nuclear protein extraction kit (Active Motif) according to manufacturer’s specifications and protein concentration was determined using a bicinchoninic acid (BCA) protein assay (Thermo Fisher Scientific). 10 µg of nuclear extract was assayed and DNA binding activities of transcription factors (ATF-2, c-Jun, FosB, Fra1, STAT-1, JunB, MEF-2, c-Myc, c-Fos, JunD, c-Jun, p50, p52, p65, RelB and c-Rel) were assessed using the TransAM transcription factor ELISA system (Active Motif). Transcription factor binding was read at 450 nm using a Synergy 2 microplate reader (BioTek, US) and expressed as fold-change relative to the uninfected (UI) control.

### Murine infection

Murine infections were performed under UK Home Office Project Licence PDF8402B7 in dedicated facilities at the University of Manchester. *A. fumigatus* spores were harvested and prepared as previously described (14) and following serial dilution, viable counts from administered inocula were determined by growth for 24–48 h on ACM. Mice (male CD1 outbred mice, *n*=5 per time point) were rendered leukopenic by administration of cyclophosphamide (150 mg/kg, intraperitoneal) on days -3, and -1 and a single subcutaneous dose of hydrocortisone acetate (112.5 mg/kg) administered on day -1. Mice were housed in individually vented cages, anaesthetized by halothane inhalation and infected by intranasal instillation of spore suspensions with 10^8^ spores of *A. fumigatus* ATCC46645. Mice were euthanised at 24 and 48 h post-infection to examine activation of pulmonary NF-κB signaling in host tissue. Immunohistochemistry was performed using the isotypic antibodies anti-p65 (ab16502, Abcam) and anti-RelB (ab180127, Abcam) and images were acquired on a 3D-Histech Pannoramic-250 microscope slide-scanner using a 20x objective (Zeiss). Snapshots of the slide-scans were taken using the Case Viewer software (3D-Histech). NF-κ? subunits (RelB and p65) immunoreactivity was assessed blindly and independently from two investigators in order to score negative, weak, moderate and strong intensity.

### Statistical analysis of data

GraphPad Prism was used to interpret data and *p* values were calculated through unpaired t tests or 1-way ordinary ANOVA tests with multiple comparison analysis as indicated. Error bars show the Standard Deviation (SD). ^****^p ≤ 0.0001, ^***^p ≤ 0.001, ^**^p ≤ 0.01, and ^*^p ≤ 0.05

## Results

### *Aspergillus fumigatus* spores, hyphae and secreted products drive epithelial decay via temporally and mechanistically distinct processes

*A. fumigatus* causes epithelial detachment and damage via multiple, temporally distinct mechanisms (3). To define the dynamics of the underlying processes and address the role of host cell activities in driving epithelial damage, A549 cells were challenged with live *A. fumigatus* spores, or culture filtrates (CF). Widefield fluorescence microscopy was used to study the gross appearance of epithelial monolayers at set time-points post-infection (Fig. 1A) and to quantify the degree of AEC detachment (Figs. 1B - 1D and S1B).

**Figure 1:**
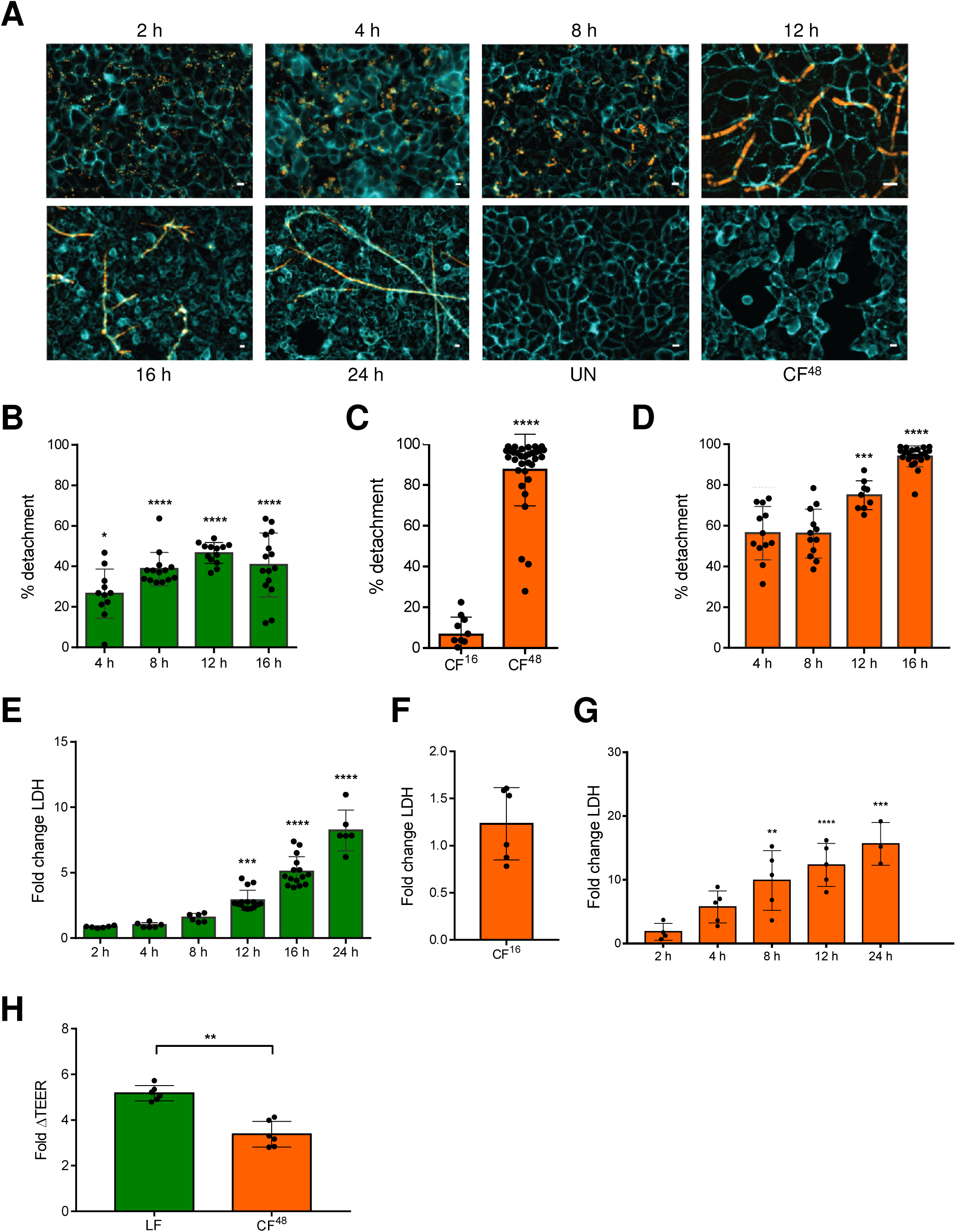
Temporal quantitative analysis of epithelial decay following challenge with live *A. fumigatus* spores or CF. **(A)** Visualisation of concanavalin A-FITC labelled A549 monolayers incubated with the *A. fumigatus* tdTomato^ATCC4664^ strain and a 5-fold diluted culture filtrate thereof (CF^48^). **(B)** Percentage of detachment of A549 cells following infection with 10^6^ *A. fumigatus* CEA10 spores at indicated time points. **(C)** Percentage of detachment of A549 cells following 16 h challenge with a 5-fold diluted filtrate from CEA10 fungal cultures (inoculum of 10^6^ spores/ml) grown for 16 h (CF^16^) or 48 h (CF^48^). **(D)** Percentage of detachment of A549 cells following challenge with a 5-fold diluted filtrate from CEA10 fungal cultures at indicated time points. **(E)** Fold change LDH release (relative to PBS challenge) at indicated time points with 10^6^ *A. fumigatus* CEA10 spores. **(F)** Fold change LDH release (relative to PBS challenge) at 24 h post-exposure to CF^16^. **(G)** Fold change LDH release (relative to PBS challenge) at indicated time points with a 5-fold diluted filtrate from CEA10 fungal cultures. **(H)** Fold change decrease in TEER between A549 monolayers incubated for 24 and 0 h with *A. fumigatus* CEA10 spores and a 5-fold diluted filtrate from CEA10 fungal cultures. Data represent three biological replicates with 1-5 technical replicates. Error bars show ± SEM. Data were analysed by non-parametric one-way ANOVA (Kruskal-wallis test) with Dunn’s multiple comparisons. Significance was calculated relative to challenge with vehicle control (PBS) unless otherwise shown by brackets. ^****^p ≤ 0.0001, ^***^p ≤ 0.001, ^**^p ≤ 0.01, and ^*^p ≤ 0.05

Germination of *A. fumigatus* was observed at approximately 8 h, followed by a phase of rapid hyphal extension between 8 and 12 h post-infection (Fig. 1A). In response to challenge with live *A. fumigatus* spores, significant detachment (∼20%) of A549 cells was detected by 4 h (Fig. 1B), when the fungus is still a spore (Figs. 1A and 1B). Upon spore germination (8 – 12 h), pneumocyte detachment further increases to reach 40-50 % (Fig. 1B). To assess the role of secreted fungal products in driving epithelial decay, AEC detachment was quantified following exposure for 16 h to CF harvested from fungal broth culture at 16 (CF^16^) or 48 (CF^48^) h. Pneumocyte detachment was caused by exposure to CF^48^ but not to CF^16^. Therefore, the cytolytic capacity of *A. fumigatus* increases >80 fold within the 16 – 48 h time frame (Fig. 1C). In contrast to live fungal spores (Fig. 1B), exposure to CF^48^ led to much more rapid epithelial decay (Fig. 1D). The magnitude of epithelial disintegration in response to challenge with spores and CF^48^ was independent of the *A. fumigatus* strain tested (Fig. S1B).

To determine whether epithelial detachment was caused by cytolytic death of AECs, lactate dehydrogenase (LDH) release was quantified over a similar time course of fungal challenge. In response to challenge with live *A. fumigatus* spores, LDH release first significantly exceeded that due to PBS challenge at 12 h post-infection (Fig. 1E), which correlated with the onset of hyphal development (Fig. 1A). LDH activity increased in a dose-dependent manner proportionate to increasing spore density of the fungal inoculum (Fig. S1C). In contrast to spore exposure where cytolytic damage takes 12 h to become detectable, AECs challenged with CF^48^ exhibited a rapid time- and dose-dependent lysis, which commenced at 2 h post-challenge (Fig. 1G and S1D). Notably, as with epithelial detachment, CF^16^ did not induce LDH release (Fig. 1F). The magnitude of epithelial cell lysis in response to challenge with spores and CF^48^ was independent of the *A. fumigatus* strain tested although timescales differed slightly (Fig. S1E).

To assess the relative contributions of hyphae versus soluble factors in driving epithelial decay, the integrity of Calu-3 monolayers was measured by monitoring changes in transepithelial resistance (TEER) at 24 h following challenge with live *A. fumigatus* spores or CF^48^ (Figs. 1H and S1F). Infection of monolayers with live fungal spores for 24 h reduced TEER by 5 fold relative to an uninfected monolayer (Fig. 1H) whereas exposure to CF^48^ for 24 h reduced TEER by only 2 to 3 fold relative to an uninfected monolayer (Fig. 1H). The decrease in TEER in response to challenge with spores and CF^48^ was independent of the *A. fumigatus* strain tested (Fig. S1G). Taken together these findings suggest that fungal spores are involved in initiating epithelial damage, which is then further propagated by i) physical invasion of the epithelial layer and ii) effector-mediated lysis of epithelial cells.

### Cytokine and extracellular matrix responses to *A. fumigatus* spores are subsequently remodelled by secreted fungal products

In addition to cytolytic damage resulting from direct host-pathogen interaction, the subversion of innate host defences can indirectly drive tissue damage via disablement or moderation of the host immune response. In order to characterise the effects of fungal challenge on host-derived cytokine and extracellular matrix (ECM) responses, AECs were exposed to live *A. fumigatus* spores or fungal secreted products (CF^48^) and expression of cytokines and (ECM) components were quantifed via (i) an unbiased high throughput immunoblotting approach (Supplementary Data 4) and (ii) targeted quantitation of nine host proteins exhibiting significant changes including IL-6 and IL-8, which have been previously reported to be induced by *A. fumigatus* in bronchial and alveolar epithelial cells (15, 16) and GM-CSF, G-CSF, IL-6 and IL-8 by *C. albicans* in a buccal epithelial carcinoma cell line (5) (Fig. 2).

**Figure 2:**
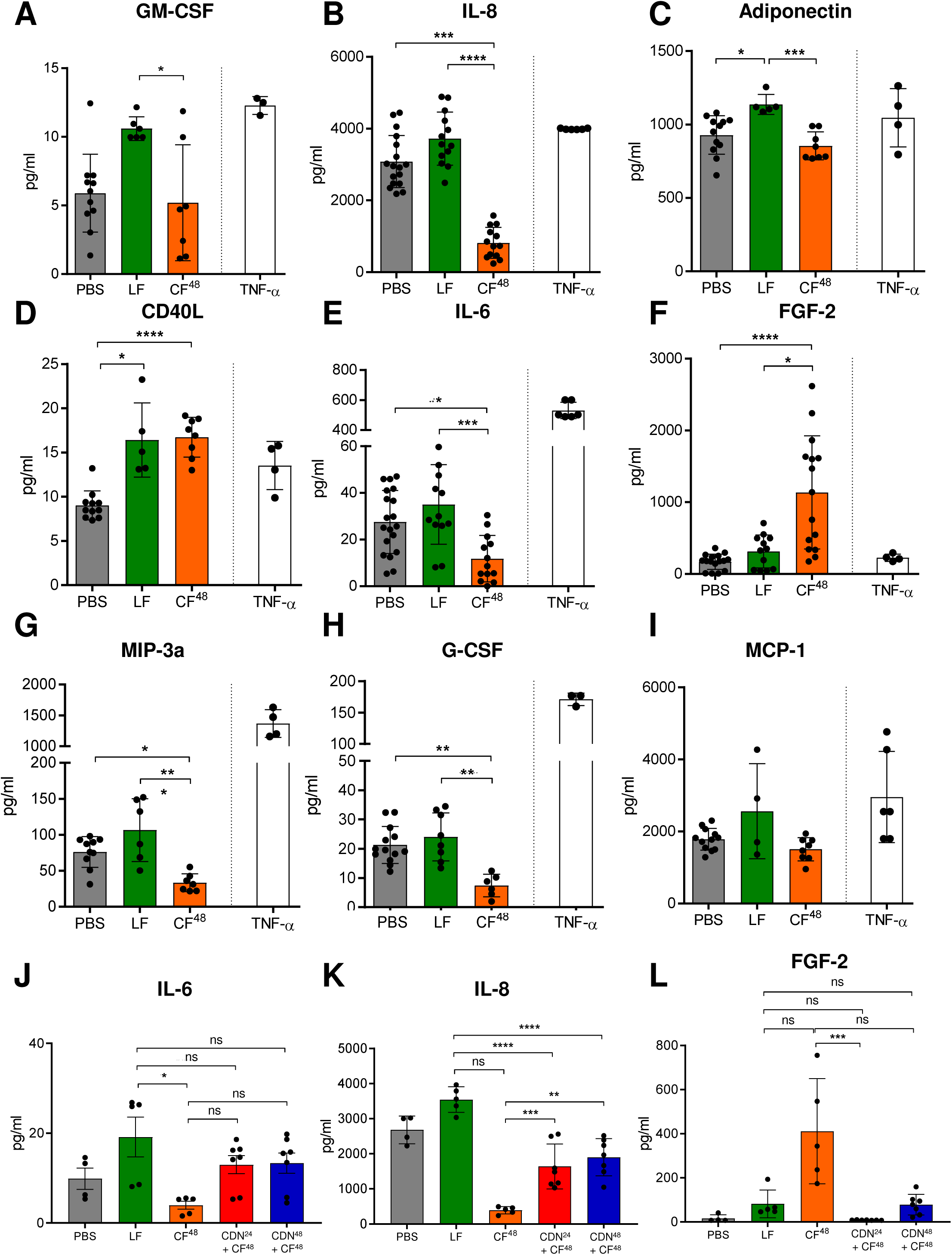
Differential cytokine secretion by A549 cells in response to *A. fumigatus* spores and CF^48^. **(A-I)** Secreted cytokines (**A**: GM-CSF; **B**: IL-8; **C**: Adiponectin, **D**: CD40L; **E**: IL-6; **F**: FGF-2; **G**: MIP-3a; **H**: G-CSF and **I**: MCP-1) were quantified in cell culture supernatant following exposure to *A. fumigatus* spores (1×10^7^ spores/ml) or 5-fold diluted CF^48^ for 24 h. **(J-L)** Cytokines (**J**: IL-6; **K**: IL-8 and **L**: FGF-2) were quantified in A549-free culture supernatants collected after exposure of epithelial monolayers to *A. fumigatus* for 24 h (CF^24^) followed by incubation with CF^48^ or with culture filtrates recovered from a 24 h infection of AECs with CF^48^ (CDN). Data represent three biological replicates with 1-5 technical replicates. Error bars show ± SEM. Data were analysed by non-parametric one-way ANOVA (Kruskal-wallis test) with Dunn’s multiple comparisons. Significance was calculated relative to challenge with vehicle control (PBS) and between each treatment as shown. ^****^p ≤ 0.0001, ^***^p ≤ 0.001, ^**^p ≤ 0.01, and ^*^p ≤ 0.05

AEC challenge with *A. fumigatus* spores for 24 h significantly increased the secretion of GM-CFS, IL-8, Adiponectin and CD40L, relative to the vehicle control (Figs. 2A, B, C and D). A non-significant increase in IL-6, FGF-2 and MIP-3a secretion was also observed (Figs. 2E, F and G). In stark contrast, relative to challenge with live fungus, challenge with CF^48^ for 24 h reduced the concentrations of most cytokines to less than basal (PBS) levels. However, secretion of FGF-2 and CD40L was significantly increased in response to CF^48^ challenge (Figs. 2D and 2F). The reductions in IL-6 and IL-8 secretion might have derived from rapid epithelial decay, or direct action of fungal proteases that degrade cytokines. In order to distinguish between the two possibilities, conditioned media harvested from epithelia culture at 24 h post-infection with live fungus (CDN^24^), and therefore exhibiting heightened cytokine concentrations, was subsequently exposed to culture filtrates obtained from mature fungal cultures grown in the presence (CDN^48^) or absence (CF^48^) of host cells. This revealed that IL-6 depletion in response to CDN^24^ challenge does not significantly differ from challenge with live fungus, thereby negating a role for fungal proteases in IL-6 degradation. Conversely, IL-8 is significantly depleted by exposure to CDN^24^ causing a ∼50% reduction in IL-8 concentration relative to challenge with live fungus (Figs. 2J-K). Epithelial challenge with CF^48^ prompted significantly increased FGF-2 secretion, an effect that could not be recapitulated via CDN^24^ challenge (Figs. 2F and 2L), thereby indicating that FGF-2 induction depends on the presence of one or multiple soluble factors produced by mature fungal hyphae. Taken together, these data indicate that *A. fumigatus* secreted products act in a target-specific manner to remodel the local cytokine and extracellular matrix environment during epithelial infection.

### Spore germination in proximity to pneumocytes iteratively stimulates NF-kB and MAPK signalling and DNA binding of downstream transcription factors

Although the ability of AECs to orchestrate immunological responses in response to *A. fumigatus* has been previously documented (16-19), the fungal morphotypes propogating intracellular signalling have remained obscure. The phosphorylation status of three mitogen-activated protein kinases (MAPKs) p38, ERK, and JNK, the Inhibitor of kappa-light-chain-enhancer of activated B cells (IκBα) and of protein kinase B (AKT), along with binding activities of downstream transcription factors (p52, p50, RelB, p65, MEF-1, c-Myc, JunD) including several that have been previously documented as responsive to *C. albicans* challenge of oral epithelial cells or keratinocytes (5, 18, 20, 21), were measured over a time-series of co-incubation with live *A. fumigatus* spores or CF^48^. The earliest evidence of a host response to fungal challenge was observed from 4 h post-infection at which time a modest, but significant, increase in IκB-α phosphorylation (Fig. 3A) was observed that persisted throughout the course of infection. A similarly modest phosphorylation of JNK (Fig. 3B) was observed. Phosphorylation of p38 (Fig. 3C), p-Akt and p-ERK1/2 (Fig. S2E) did not occur in response to challenge with spores.

**Figure 3:**
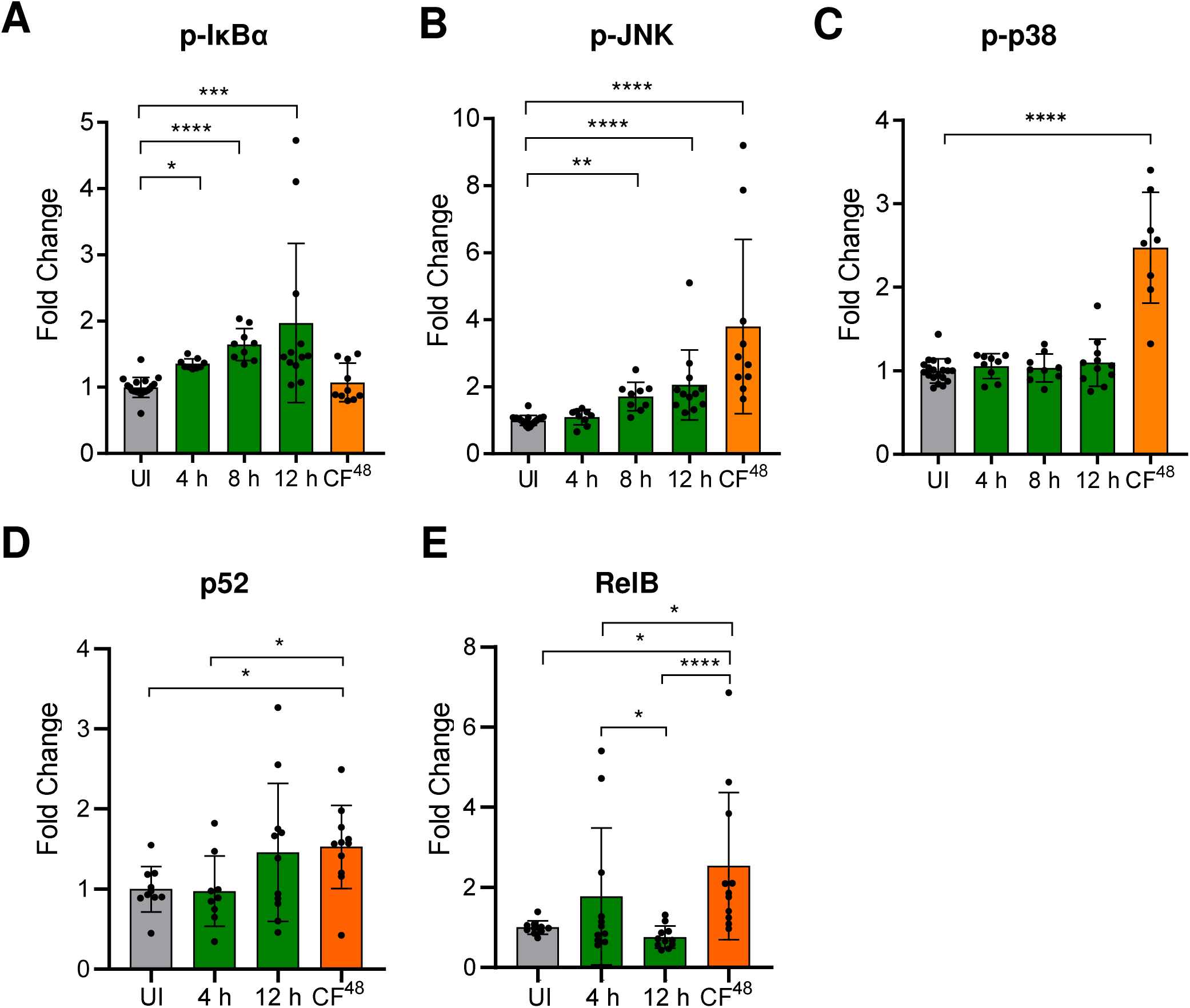
Different fungal morphotypes induce differential and dynamic host responses in A549 cells. **(A-C)** Fold change (relative to uninfected control, UI) phosphorylation of NF-kB (**A**: p-IkBα) and MAPK (**B**: p-JNK and **C**: p-p38) signalling following exposure to *A. fumigatus* spores (1×10^7^ spores/ml) for indicated time points or 5-fold diluted CF^48^ for 4 h. **(D-E)** Fold change (relative to uninfected control, UI) in DNA binding activity of non-canonical NF-kB transcription factors (**D**: p52 and **E**: RelB) following exposure to *A. fumigatus* spores (10^7^ spores/ml) for indicated time points or 5-fold diluted CF^48^ (inoculum of 10^6^ spores/ml) for 4 h. Data represent three biological replicates with 1-5 technical replicates. Error bars show ± SEM. Data was analysed by non-parametric one-way ANOVA (Kruskal-wallis test) with Dunn’s multiple comparisons. Significance was calculated relative to challenge with vehicle control (PBS) as shown. ^****^p ≤ 0.0001, ^***^p ≤ 0.001, ^**^p ≤ 0.01, and ^*^p ≤ 0.05

Exposure of A549 cells to CF^48^ led to an immediate increase in phosphorylation (3 and 1.5 fold respectively, p<0.0001) of JNK and p38 at 4 h post-exposure (Figs. 3B and 3C). In contrast, phosphorylation of IkBα, a key event in NF-kB signalling activation, was not observed during challenge with CF^48^ (Fig. 3A), suggesting that NF-kB activation in response to *A. fumigatus* challenge is most likely contact-mediated. The magnitude of IκB-α, JNK and p38 phosphorylation in response to challenge with spores was independent of the *A. fumigatus* strain tested (Figs. S2A-C). However, in line with an expanding number of studies showing strain to strain variation of *A. fumigatus* phenotypes, immune responses and virulence in various models of infection (22), CF^48^ from the three parental isolates tested (CEA10, ATCC46645 and a non-homologous end joining deficient mutant *ΔakuB*^KU80^) elicited JNK and p38 phosphorylation to significantly different extents (Figs. S2A-C). In our experimental setup, no changes in Akt or ERK phosphorylation were observable following challenge with CF^48^ (Figs. S2D-E).

Activation of distinct signalling mechanisms by live spore challenge and CF^48^ was paralleled by differential modulation of DNA binding of associated transcription factors (Figs. 3D-E and S4). Upon epithelial challenge with live *A. fumigatus* for 12 h, DNA binding of Jun D and MEF-2 decreased significantly (0.5 fold, p<0.05 and p<0.01 respectively), while DNA binding of c-Myc increased (1.75 fold, p<0.05) (Figs. S3A-C). Importantly, CF^48^ challenge of epithelial monolayers for 4 h led to a significant increase in DNA binding (1 and 3 fold, respectively, p<0.05) of the p52 and RelB components of non-canonical NF-κB signalling (Fig. 3D-E). No significant changes in ATF-2, c-Jun, FosB, c-Fos, Fra1, c-Rel, STAT-1, JunB (not shown), p50 (Fig. S3D), or p65 (Fig. S3E) binding activity were observed in our population-level assays. However, extensive variation of the experimental readings for these transcription factors was observed, suggesting rapidly cycling modulation of their activity in response to fungal challenge.

### *A. fumigatus*-induced epithelial damage is mediated by IκB-α and p-JNK activation

Epithelial perturbation, detachment and lysis are key features of *A. fumigatus in vitro* infections first in a contact-dependent manner and subsequently via the secretion of fungal soluble factors (Fig. 1) (3). Notably, peak phosphorylation of both IκB-α and JNK is observed at 12 h post-infection with live fungus (Fig. 3A-B), thereby suggesting that hyphal-dependent activation of the NF-kB and MAPK pathways leads to *A. fumigatus* induced epithelial damage. In order to determine if phosphorylation of IκB-α and JNK requires live fungus, *A. fumigatus* was grown for 12 h to induce hyphae formation and A549 AECs were challenged with pre-grown heat-killed and live *A. fumigatus*. Contrary to live *A. fumigatus* pre-grown as hyphae (up to 1.75 fold increase for IκB-α and JNK relative to uninfected, p ≤ 0.001), the heat-killed morphotypes failed to induce p-IκB-α and p-JNK activation upon challenge of epithelial monolayers, indicating that fungal viability is a pre-requisite for phosphorylation of IκB-α and JNK (Figs. 4A-B).

**Figure 4:**
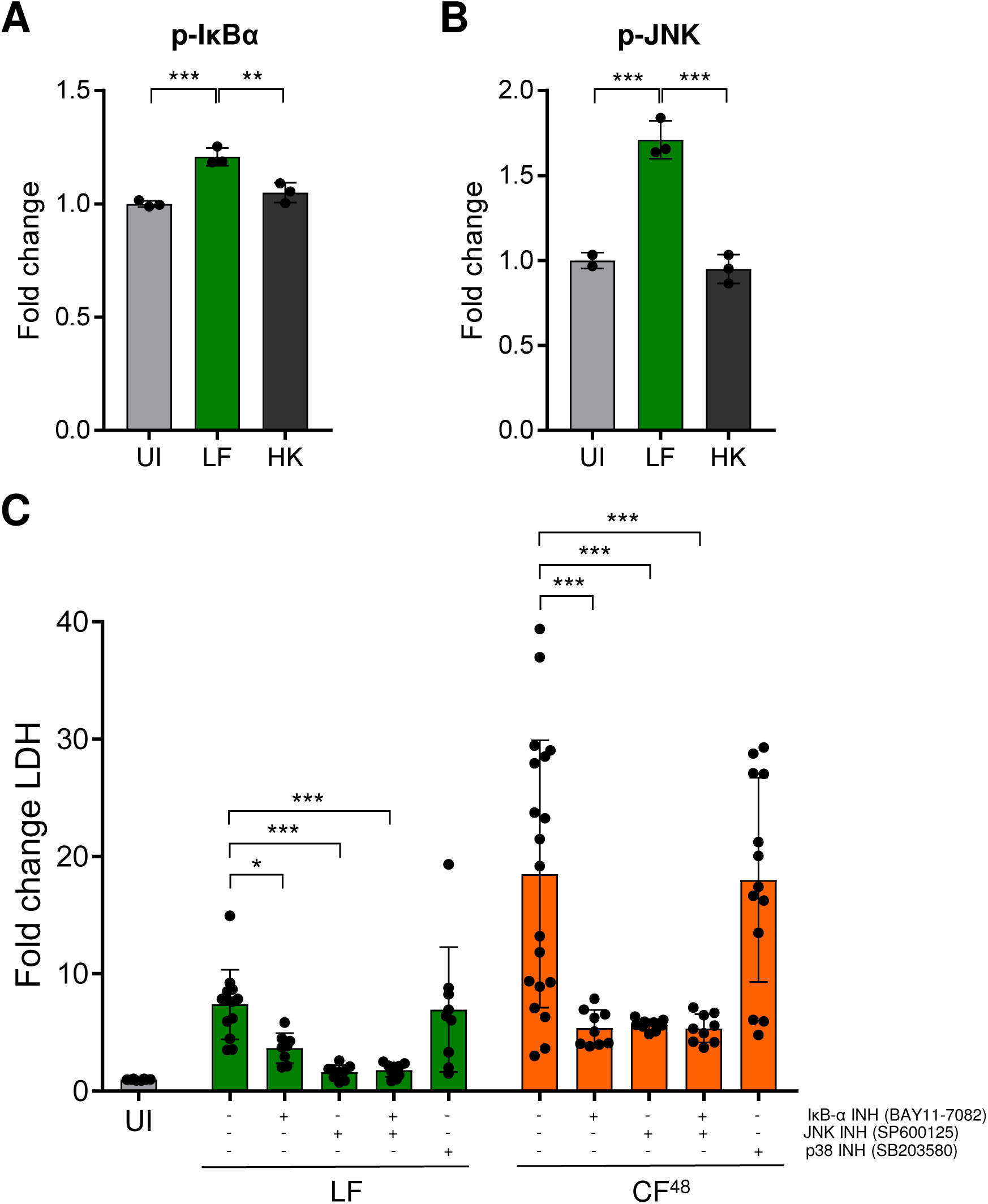
Signalling activation is dependent on fungal viability and plays a role in epithelial damage. **(A-B)** Fold change (relative to uninfected control, UI) phosphorylation of NF-kB (**A**: p-IkBα) and MAPK (**B**: p-JNK) signalling following 4 h exposure to live and heat killed morphotypes (10^7^ spores/ml) of *A. fumigatus* pre-grown to hyphae for 12 h. **(C)** Fold change LDH release (relative to PBS challenge) following chemical inhibition of NF-kB (IkBα inhibithor BAY11-7082) and MAPK (JNK inhibitor SP600125 and p38 inhibitor SB203580) signalling pathways and exposure to *A. fumigatus* spores (10^6^ spores/ml) or 5-fold diluted CF^48^ for 24 h. Data represent three biological replicates with 1-5 technical replicates. Error bars show ± SEM. Data were analysed by non-parametric one-way ANOVA (Kruskal-wallis test) with Dunn’s multiple comparisons. Significance was calculated relative to challenge with vehicle control (PBS) and between each treatment as shown. ^****^p ≤ 0.0001, ^***^p ≤ 0.001, ^**^p ≤ 0.01, and ^*^p ≤ 0.05

The relative contribution of NF-κB and MAPK pathways in AEC lysis was determined using small chemical inhibitors against IκB-α (BAY11-7082), JNK (SP600125) and p38 (SB203580) (Fig. 4C). JNK inhibition significantly reduced hyphal- and CF^48^-induced epithelial lysis by over 60%. Compared to JNK inhibition, IκB-α inhibition yielded a less pronounced protective effect for hyphal-induced lysis, but similar reduction in CF^48^ induced AEC lysis. No evidence of an additive or synergistic effect was observed when both IκB-α and JNK pathways were inhibited simultaneously. Moreover, p38 inhibition did not impact epithelial damage upon challenge with live fungus or CF^48^. Taken together, these results indicate that *A. fumigatus*-induced epithelial damage is mediated by IκB-α and JNK, but not p38 activation. ’

### JNK-mediated epithelial lysis requires *A. fumigatus* PacC, GliP and PrtT

Functional genomic studies have demonstrated that the pH sensing regulator PacC governs *in vitro* epithelial damage and *A. fumigatus* pathogenicity in leukopenic mice, by modulating expression of over 250 secreted proteins, in addition to the secondary metabolite gliotoxin and cell wall associated proteins (3). However, deletion alone of the conserved positive regulator of secreted proteases PrtT (23, 24) or of the nonribosomal peptide synthetase GliP, essential for gliotoxin production (25-27), does not diminish survival of leukopenic mice following *A. fumigatus* challenge. Taken together, these findings suggest redundancy and possibly cooperative action of soluble effectors such as proteases and secondary metabolites in driving epithelial damage and *A. fumigatus* pathogenicity.

To determine the role of PacC, gliotoxin biosynthesis and protease production in promoting epithelial detachment (Fig. 5A), lysis (Fig. 5B) and immune responses (Fig. 6), AEC monolayers were infected with an isogenic panel of *A. fumigatus ΔpacC, ΔgliP* and *ΔprtT* mutants and respective CF^48^. In agreement with data previously published by Bertuzzi *et al*., 2014 (3), both live spores and CF^48^ of the *ΔpacC* mutant induce significantly less epithelial cell detachment (30% and 75% respectively) than the parental isolate (Fig. 5A). In contrast to *ΔpacC* challenges, CF^48^ but not live spores of the *ΔprtT* and *ΔgliP* mutants showed a decreased capacity to elicit epithelial cell detachment compared to the respective parental isolate (50% reduction, p ≤ 0.0001), while live fungal spores of the two mutants induced a similar level of cell detachment compared to their parental counterpart (Fig. 5A). Importantly, the attenuation of epithelial cell detachment caused by *ΔprtT* and *ΔgliP* CF^48^ was significantly less pronounced than the attenuation measured for *ΔpacC* CF^48^ (p ≤ 0.0001) (Fig. 5A), thereby supporting additivity or cooperativity of secreted proteases and gliotoxin in eliciting epithelial cell detachment at early time-points of *in vitro* infection.

**Figure 5:**
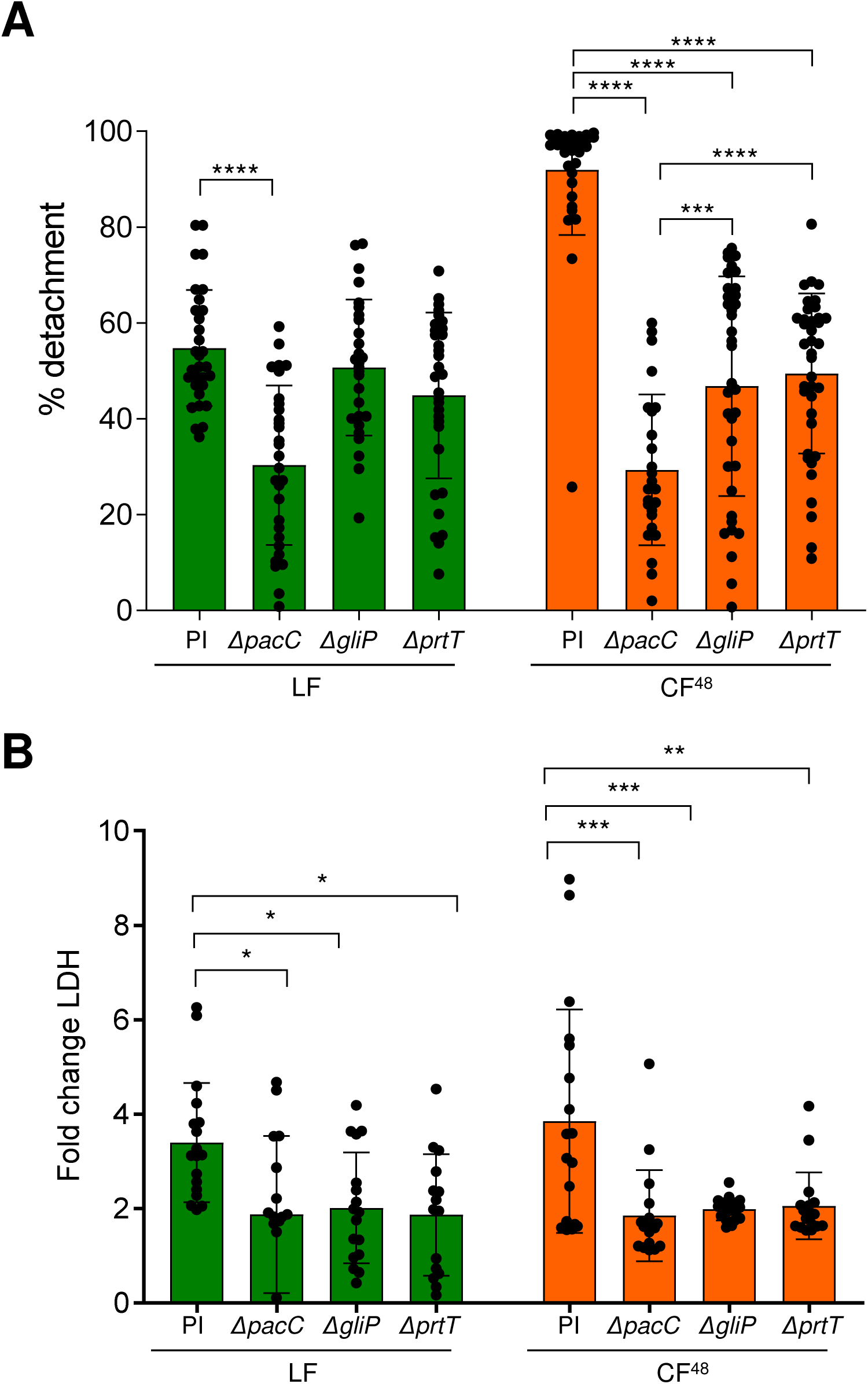
*A. fumigatus* Δ*pacC*, Δ*gliP* and Δ*prtT* mutants and respective CF^48^ fail to induce AEC detachment and lysis. **(A)** Percentage of detachment of A549 cells following infection with 10^6^ *A. fumigatus ΔpacC, ΔgliP* and *ΔprtT* mutants and respective parental isolate (PI) and CF^48^ thereof for 16 h. **(B)** Fold change LDH release (relative to PBS challenge) following infection with 10^6^ *A. fumigatus ΔpacC, ΔgliP* and *ΔprtT* mutants and respective parental isolate (PI) and CF^48^ thereof for 24 hours. Data represent three biological replicates with 1-5 technical replicates. Error bars show ± SEM. Data were analysed by non-parametric one-way ANOVA (Kruskal-wallis test) with Dunn’s multiple comparisons. Significance was calculated relative to the parental isolate (PI) as shown. ^****^p ≤ 0.0001, ^***^p ≤ 0.001, ^**^p ≤ 0.01, and ^*^p ≤ 0.05

**Figure 6:**
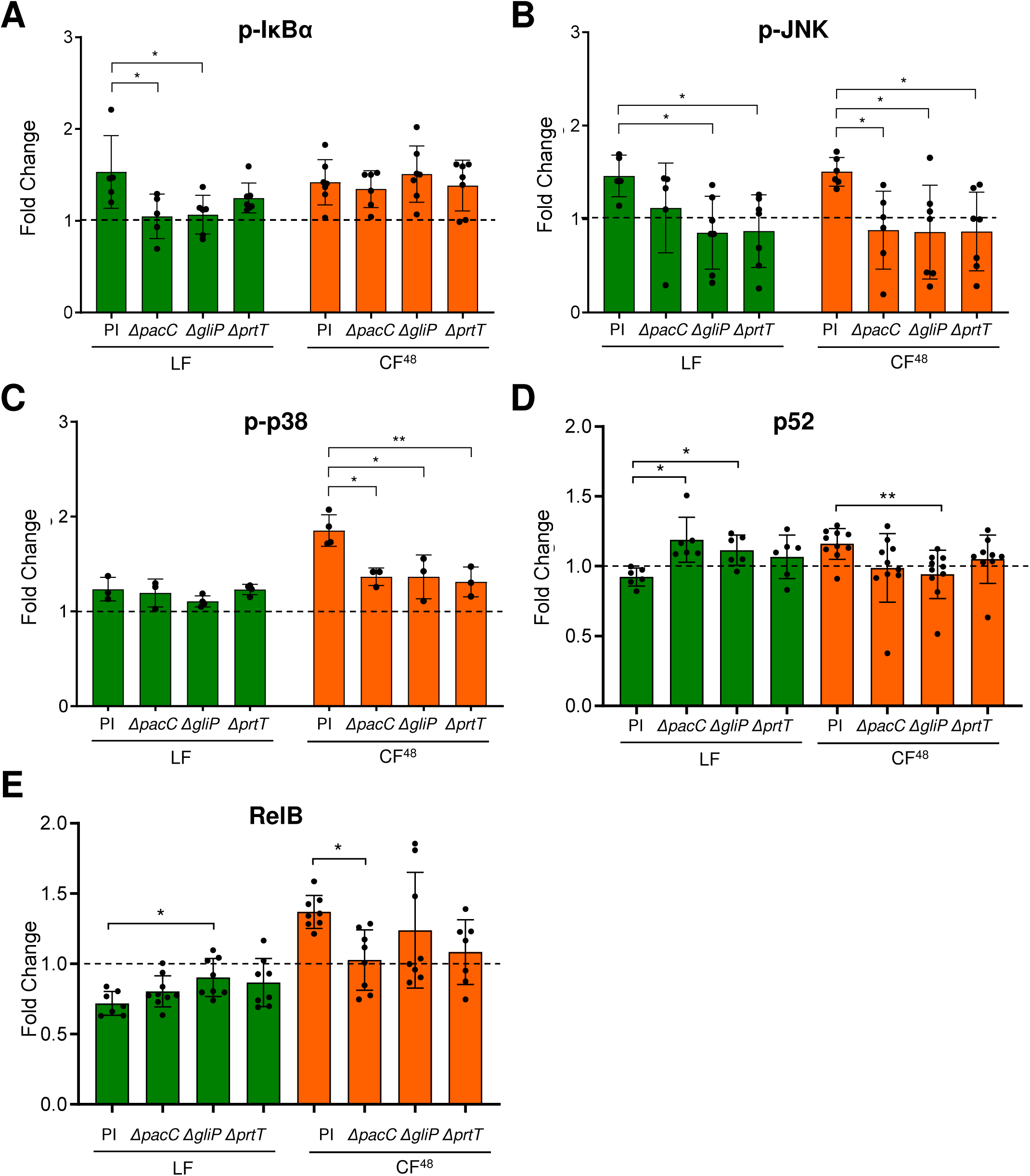
*A. fumigatus* Δ*pacC*, Δ*gliP* and Δ*prtT* mutants and respective CF^48^ fail to activate AEC host signalling proteins and transcription factors. **(A-C)** Fold change (relative to PBS challenge) phosphorylation of NF-kB (**A**: p-IkBα) and MAPK (**B**: p-JNK and **C**: p-p38) signalling following exposure to *A. fumigatus ΔpacC, ΔgliP* and *ΔprtT* mutants and respective parental isolate (PI) (1×10^7^ spores/ml) for 8 h or respective 5-fold diluted CF^48^ for 4 h. **(D-E)** Fold change (relative to PBS challenge) in DNA binding activity of non-canonical NF-kB transcription factors (**D**: p52 and **E**: RelB) following exposure to *A. fumigatus ΔpacC, ΔgliP* and *ΔprtT* mutants and respective parental isolate (PI) (1×10^7^ spores/ml) for 8 h or respective 5-fold diluted CF^48^ for 4 h. Data represent three biological replicates with 1-5 technical replicates. Error bars show ± SEM. Data was analysed by non-parametric one-way ANOVA (Kruskal-wallis test) with Dunn’s multiple comparisons. Significance was calculated relative to the parental isolate (PI) as shown. ^****^p ≤ 0.0001, ^***^p ≤ 0.001, ^**^p ≤ 0.01, and ^*^p ≤ 0.05

In contrast to the respective parental isolate, *ΔpacC, ΔgliP* and *ΔprtT* mutants failed to induce epithelial cell lysis (Fig. 5B). Accordingly, LDH release in response to challenge with *ΔpacC, ΔgliP* and *ΔprtT* live mutants and respective CF^48^ was equivalent to the levels of the uninfected control and significantly reduced compared to challenges with the respective parental isolate (2.5 and 3 fold reduction for live fungus, p ≤ 0.05, and CF^48^, p ≤ 0.001, respectively).

Redundancy in GliP and PrtT-mediated effects on epithelial responses was also observed at the level of signalling and transcriptional activation (Fig. 6). In comparison with live fungal challenge with their respective parental isolate, activation of IκBα phosphorylation was significantly reduced upon challenge of monolayers with live *A. fumigatus* mutants lacking PacC, GliP or PrtT (Fig. 6A). However, while the *ΔprtT* mutant showed only a moderate decrease in the activation of IκBα phosphorylation after 8 h of *in vitro* infection with live spores, the *ΔpacC* and *ΔgliP* mutants completely failed (p ≤ 0.05 relative to parental isolate challenge) to elicit IκBα activation (Fig. 6A). Similarly to CF^48^ challenge with the parental strain, none of the mutants induced IκBα phosphorylation in epithelial cells upon challenge with the respective CF^48^ (Fig. 6A). While both live parental isolate and respective CF^48^ were able to induce JNK phosphorylation, *ΔpacC, ΔgliP* and *ΔprtT* mutants and CF^48^ thereof failed (p ≤ 0.05 relative to parental isolate challenges) to activate JNK (Fig. 6B). As opposed to *ΔgliP* and *ΔprtT*, however, challenges with a live *ΔpacC* isolate only resulted in a moderate reduction in JNK activation after 8 h of *in vitro* infection (Fig. 6B). Neither the parental isolate nor the mutants induced p38 phosphorylation in epithelial cells upon challenge with live fungus but, contrary to CF^48^ challenge with the parental strain, CF^48^ from the *ΔpacC, ΔgliP* and *ΔprtT* mutants failed to significantly elicit p38 activation (Fig. 6.C).

Relative to parental isolate challenges, an increase in the DNA binding of p52 and RelB was observed in response to live fungal challenges with the *ΔpacC, ΔgliP* and *ΔprtT* mutants (Figs. 6D-E). However, the same increase in the DNA binding of p52 and RelB was not detected when comparing challenges with CF^48^ of the parental isolate and the *ΔpacC, ΔgliP* and *ΔprtT* mutants. Taken together, these results highlight that PacC-mediated transcriptional regulation, gliotoxin biosynthesis and PrtT-mediated protease production critically collaborate to differentially induce MAPK and NF-κB mediated signalling in epithelial cells in response to challenge with live spores and CF^48^, therefore ultimately impacting on epithelial detachment, lysis and responses during *A. fumigatus* infections.

### Spore inhalation by leukopenic mice subverts NF-κB homeostasis

Exposure of epithelial monolayers to *A. fumigatus in vitro* leads to the iterative activation of canonical and non-canonical NF-kB signalling and respective transcriptional events as shown by the significant phosphorylation of IκB-α at 4 to 12 h post-infection with live fungus (Fig. 3A) and by the significant increase in DNA binding of the p52 and RelB components of the NF-κB signalling pathway following CF^48^ challenge (Fig. 3D-E). Importantly, *in vitro*, activation of NF-kB signalling leads to *A. fumigatus* induced epithelial damage, as demonstrated by the significant reduction of LDH release following IκB-α phosphorylation and proteolysis in monolayers challenged with live fungus and respective CF^48^ (Fig. 4C). Furthermore, the non-invasive and attenuated A. *fumigatus* mutant lacking the transcription factor PacC evokes muted canonical and non-canonical NF-κB activity compared to that induced by an isogenic parental isolate (Figs. 5 and 6), strongly implicating NF-κB in damage-inducing host responses to fungal challenge during infection.

Aberrant activation or disruption of NF-kB signalling, in particular via the RelB-p52 pathway has been implicated in modulation of lung pathology. For example, activation of both classical and alternative NF-κB pathways has been shown to occur within the airway epithelium and may coordinately contribute to allergic inflammation airway remodeling in response to house dust mite challenge (28) and lung tissues from patients with acute respiratory distress syndrome (ARDS) or pulmonary adenocarcinoma exhibit increased expression of p100/p52 (29, 30). p52 over-expression in airway epithelial cells of transgenic mice causes increased lung inflammation, injury, and mortality following LPS stimulation; and increased tumor number and progression after injection of the carcinogen urethane (29, 30). Conversely, overexpression of RelB in mouse airways has been shown to dampen acute smoke-induced pulmonary inflammation (31).

In order to assess the physiological relevance of disrupted NF-κB homeostasis in an intact tissue setting, leukopenic mice were infected by intranasal instillation of *A. fumigatus* spores and euthanised at 24 and 48 h post-infection to examine activation of pulmonary NF-κB signaling in host tissue (Fig. 7). A semi-quantitative immunohistochemical approach was applied to comparatively assess nuclear translocation of the NF-κ? p65 and RelB subunits in the vicinity of infectious lesions (Fig. 7A). In uninfected tissues, at both 24 and 48 h post-infection, p65 negative nuclei (72-76%) significantly outnumbered p65 positive nuclei (Figs. 7B-C). In contrast, in infected tissues at both 24 and 48 h post-infection, equivalent numbers of p65 positive (44-39%) and negative (56-61%) nuclei were observed (Figs. 7B-C). At both time-points analysed, positivity for RelB immunoreactivity was significantly increased in infected tissues with 57-61% of nuclei scoring as RelB positive in infectious lesions compared to only 37-40% in uninfected tissue (Figs. D-E). These findings substantiate the physiological relevance of our *in vitro* findings (Fig. 3E and Fig. S3E), and confirm that dysregulation of the NF-kB signaling axis is a critical component of *A. fumigatus*-mediated lung infection.

**Figure 7:**
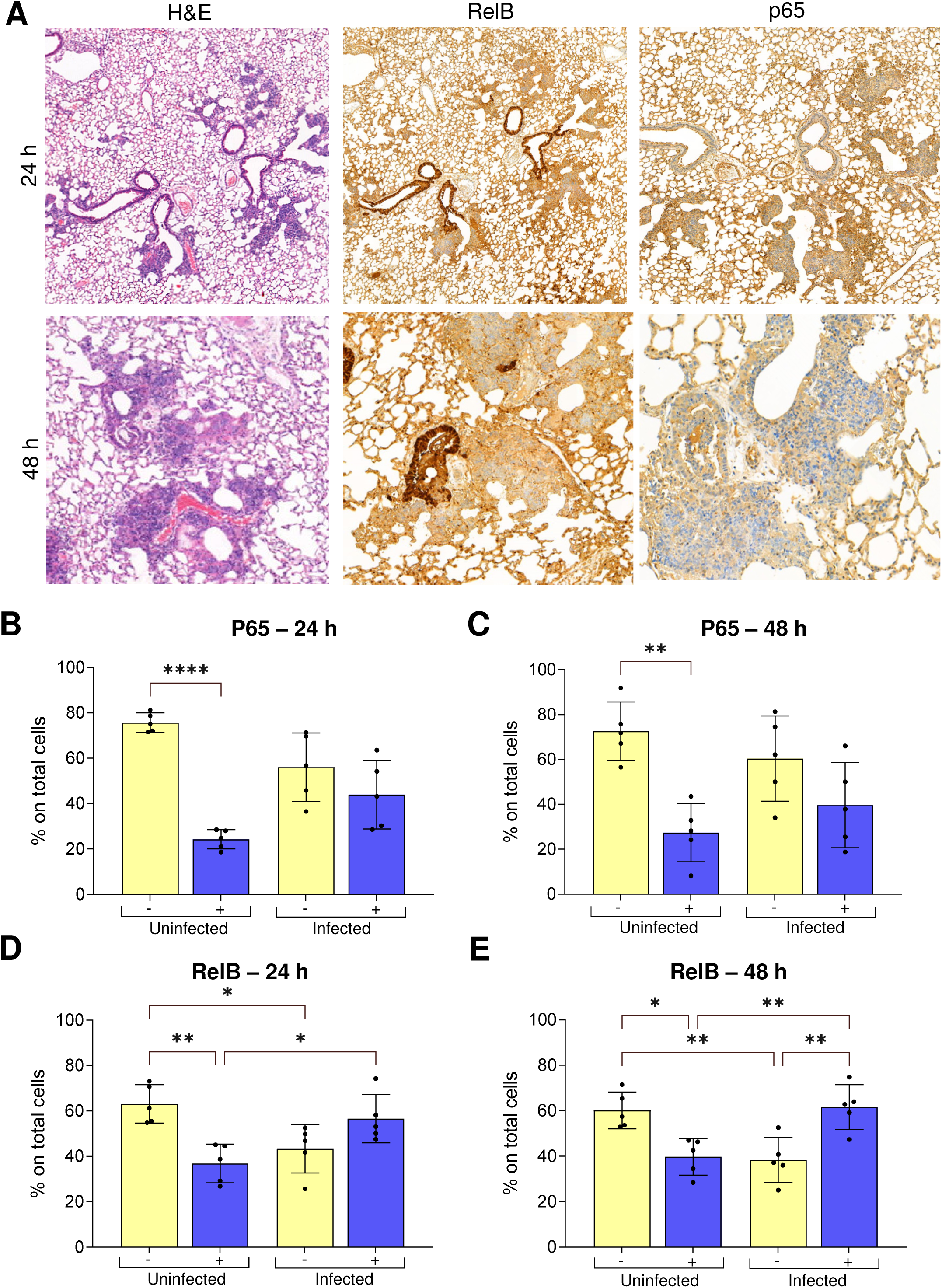
Immunohistochemistry to assess RelB and p65 immunoreactivity *in vivo* indicates iterative subvertion of the NF-κB signalling axis following *A. fumigatus* infection of leukopenic mice. **(A)** Hematoxylin and Eosin (H&E) staining and RelB/p65 immunochemistry staining of histological sections from leukopenic mice infected with *A. fumigatus* dTomato^ATCC46645^ for 24 and 48 h **(B-E)** Quantification of p65 **(B&C)** and RelB **(D&E)** staining from scoring infected lesions and uninfected tissue in histological sections from leukopenic mice. ^****^p ≤ 0.0001, ^***^p ≤ 0.001, ^**^p ≤ 0.01, and ^*^p ≤ 0.05

## Discussion

One key pathological feature of pulmonary aspergillosis is gross destruction of the lung parenchyma resulting from fungus-mediated damage, however the precise mechanism(s) by which fungal hyphae penetrate the lining of the lung are unknown. Here we show that the obligatory spore to hyphal morphological transition of *A. fumigatus* exposes epithelial cells to a wide range of cell surface-associated and secreted fungal antigens and toxins that are recognised by the host leading to activation of multiple different host signalling pathways. Previous studies showed that epithelial damage results from the combined effect of fungal secreted products (3, 32, 33) and aberrant activation of host cell signalling (18). However the available data are sparse and fragmented, and focus on single timepoints and morphotypes, thereby failing to account for the morphological dynamism in fungal growth that effects differential immunogenic responses. In this study, we sought to understand the AEC-*Aspergillus* interaction in a manner that captures the morphogenic transitions of the pathogen in a real infection by elucidating the dynamic host response to the different morphotypes of *A. fumigatus* and the contribution of each stage to epithelial damage. We found that airway epithelial cells respond in a distinctive and dynamic manner to *A. fumigatus* conidia, germlings, hyphae and secreted fungal products via (i) immediate and sustained activation of the canonical NF-κB signaling circuit and (ii) subsequent JNK and p38 activation. Since aberrant activation of inflammatory signalling pathways can impact epithelial integrity and inhibition of such signalling has a protective effect (this work and Sharon et al., 2011 (18)), we surmise that such processes could be modulated therapeutically to protect the integrity of infected lung tissue.

This study validates our earlier findings (3) that *A. fumigatus*-mediated epithelial detachment occurs via at least two distinct mechanisms; firstly in a contact-dependent manner and secondly by soluble hyphae-derived factors. We reveal a previously unappreciated degree of mechanistic complexity involving: contact-mediated perturbation, physical invasion of the epithelial stratum by *A. fumigatus* germlings and hyphae and host cell lysis via activity of soluble effectors. Our data demonstrate co-operativity between physical fungal invasion and input from soluble effectors in achieving the overall extent of damage to the host. A similar mechanism has been demonstrated for invasive *C. albicans* infections whereby physical force exerted by the growing hyphae on the epithelium is complemented by specific virulence factors including the invasin Als3, several secreted proteases such as Sap5 and the peptide toxin candidalysin, which together promote degradation of tight junction proteins and barrier/membrane disruption (34-37). Our data support the hypothesis that physical invasion by *A. fumigatus* hyphae may be important in reducing the interepithelial adhesion, thereby initiating cell detachment, whereas soluble products are required for inducing lytic cell death. We demonstrated that lytic epithelial damage is elicited via host immune responses to fungal challenge which when ablated, using small molecule inhibitors, resulted in a significant degree of protection.

We show that cultured AECs respond in a distinctive and dynamic manner to *A. fumigatus* infection during the early (conidia), intermediate (germlings/immature hyphae) and late (mature hyphae/secreted effector) phases of infection, with activation of the NF-κB signaling circuit occurring within 4 h of spore exposure (Fig. 3) followed by activation of JNK signalling as spores swell and germinate (Fig. 3). As the hyphae mature, further alterations in MAPK signaling occur involving modest activation of c-myc and concomitant decreases in MEF2 and JunD activity (Fig. S3). Finally, in the late phase, hyphal-secreted components drive an increase in both JNK and p38 signalling (Fig. 3). These responses are similar to interactions of *C. albicans* and dermatophyte species with oral or skin epithelia, respectively (20). Interestingly, though, unlike *Candida* species and dermatophytes, stimulation of A549 cells with *A. fumigatus* did not activate p38 signalling in response to direct contact (20). Our results on NF-κB mediated responses to conidia and germ-tubes are in line with the only previous study that investigated the role of NF-κB signalling in airway *Aspergillus* infection; however, only a single time point of 15 h (germlings/immature hyphae) was investigated previously (16). In our study, activation of canonical NF-κB signalling occured via a contact-dependent mechanism, since activation was absent in response to challenge with soluble hyphal products (Fig. 3). The absence of canonical NF-κB signalling following challenge with CF^48^ suggests that the ligands or effectors inducing NF-κB are not secreted proteases or soluble factors; nonetheless, the inhibition of canonical NF-κB signalling significantly reduces subsequent AEC lysis demonstrating an unfavourable outcome for host cells that subsequently mount additional responses (Fig. 4C).

Consistent with previous studies, activation of host signalling in response to early *A. fumigatus* infection resulted in significant increases in cytokine expression including IL-8 and IL-6. For the first time, we report increased expression of FGF-2(basic) release from AECs in response to early *A. fumigatus* infection. This finding is concordant with increased FGF-2(basic) protein expression reported previously in lungs of infected rats (38) and our data that demonstrate resilience of FGF-2(basic) to proteolytic degradation by secereted fungal proteases (Fig 2). Exposure of AECs to CF^48^ alone leads to significant reduction in cytokine expression and production in a concentration dependent manner, suggesting the role of soluble factors in degrading these cytokines (39-41). The degradation in cytokines is likely mediated by proteases and possibly in a target specific manner as some host-derived signalling factors, such as FGF-2 and CD40L, are resistant to degradation even following exposure to CF^48^ (Fig. 2). The likely physiological relevance of such observations is substantiated by a recent study which demonstrated the ability of the pathogenic yeast of *Blastomyces dermatitidis* to down play host immune responses by elaborating the activities of fungal dipeptidyl-peptidase IVA (DppIVA), a close mimic of the mammalian ectopeptidase CD26 (42, 43). DppIVA was shown to cleave human chemokines in a C-C (43) and C-X-C (42) targeted manner. Worthy of note is that the expression of the *dppIV* homologue in *A. fumigatus* is under PacC-mediated regulation in a murine model of invasive aspergillosis (3), a factor which might contribute to target-specific modulation of host responses and inability of PacC null mutants to invade the respiratory epithelium.

The events identified from this study as leading to AEC decay appear to be mechanistically due to a combination of factors under the *A. fumigatus* master regulator PacC. Although the cohort of PacC regulated factors mediating the various damage events likely differ, it appears that fungal proteases and gliotoxin might act additively to bring about AEC lysis (Fig. 5). Interestingly, GliP and PrtT exhibit non-redundant phenotypes with respect to both JNK phosphorylation and lytic death of AECs (Fig. 5 and 6). One explanation, supported by the PacC null phenotype, is the co-operativity of gliotoxin and secreted proteases in activating host signalling. Accordingly, neither gliotoxin nor prtT null mutants are attenuated for pathogenicity in leukopenic mice (24, 26, 44, 45), but a mutant lacking both capabilities (PacC) is non-invasive and avirulent (3).

Although this work used an *in vitro* model of alveolar carcinoma cells, the reproduction of key features in a murine infection model show that A549 cells provide a robust model to study the global and temporal basis of AEC responses to *A. fumigatus*. In keeping with our *in vitro* finding that iterative stimulation of signalling via canonical and non-canonical NF-kB pathways results from exposure of epithelia to spores and soluble fungal factors, respectively, histological analysis of infected murine tissue revealed elevated immunoreactivity of isotypic anti-p65 and anti-RelB antibodies in infected versus uninfected tissues (Fig. 7). Whilst both classical and alternative NF-kB signaling can be co-ordinately activated in murine alveolar epithelial cells in response to diverse stimuli (28), we do not find that RelB expression derives solely from prior activation of canonical signaling. Therefore, it is likely that amongst the secreted factors released by mature fungal hyphae, there exists a stimulus of RelB expression. It is not yet clear whether *A. fumigatus* spore- or CF^48^-induced RelB expression acts to dampen or worsen inflammatory responses and epithelial damage but it is conceivable that either scenario could disadvantage antifungal host defences, particularly when coupled to reorchestration of inflammatory cytokine expression and cytolytic activities concomitantly expressed during hyphal invasion of the lung.

## Conclusions

This study established an *in vitro* model of dynamic and temporal AEC responses to *A. fumigatus* infections. AECs differentially recognize the different morphotypes of *Aspergillus* by mounting distinct and effective immune responses, further reinforcing the fact that the airway epithelium is functionally crucial in anti-*Aspergillus* immunity. Furthermore, the study demonstrated that during interaction with *A. fumigatus*, the AEC responses could be either destructive or protective and, in most cases, the destructive responses are driven in response to fungal secreted factors.

## Acknowledgements

Special thanks goes to Roger Meadows for his help with the Slide-scanner. Bioimaging Facility microscopes used in this study were purchased with grants from BBSRC, Wellcome and the University of Manchester Strategic Fund.This work was supported by grants from the Medical Research Council (New Investigator Research Grant MR/V031287/1 to MBe and project grants MR/S001824/1, MR/M02010X/1 and MR/L000822/1 to EMB); the Wellcome Trust (214229_Z_18_Z to JRN and Strategic Award for Medical Mycology and Fungal Immunology 097377/Z/11/Z to JRN and EMB); the NIH Research at Guys and St. Thomas’s NHS Foundation Trust and the King’s College London Biomedical Research Centre (IS-BRC-1215-20006 to JRN).

## Supplementary Figures

**Supplementary Figure 1:**
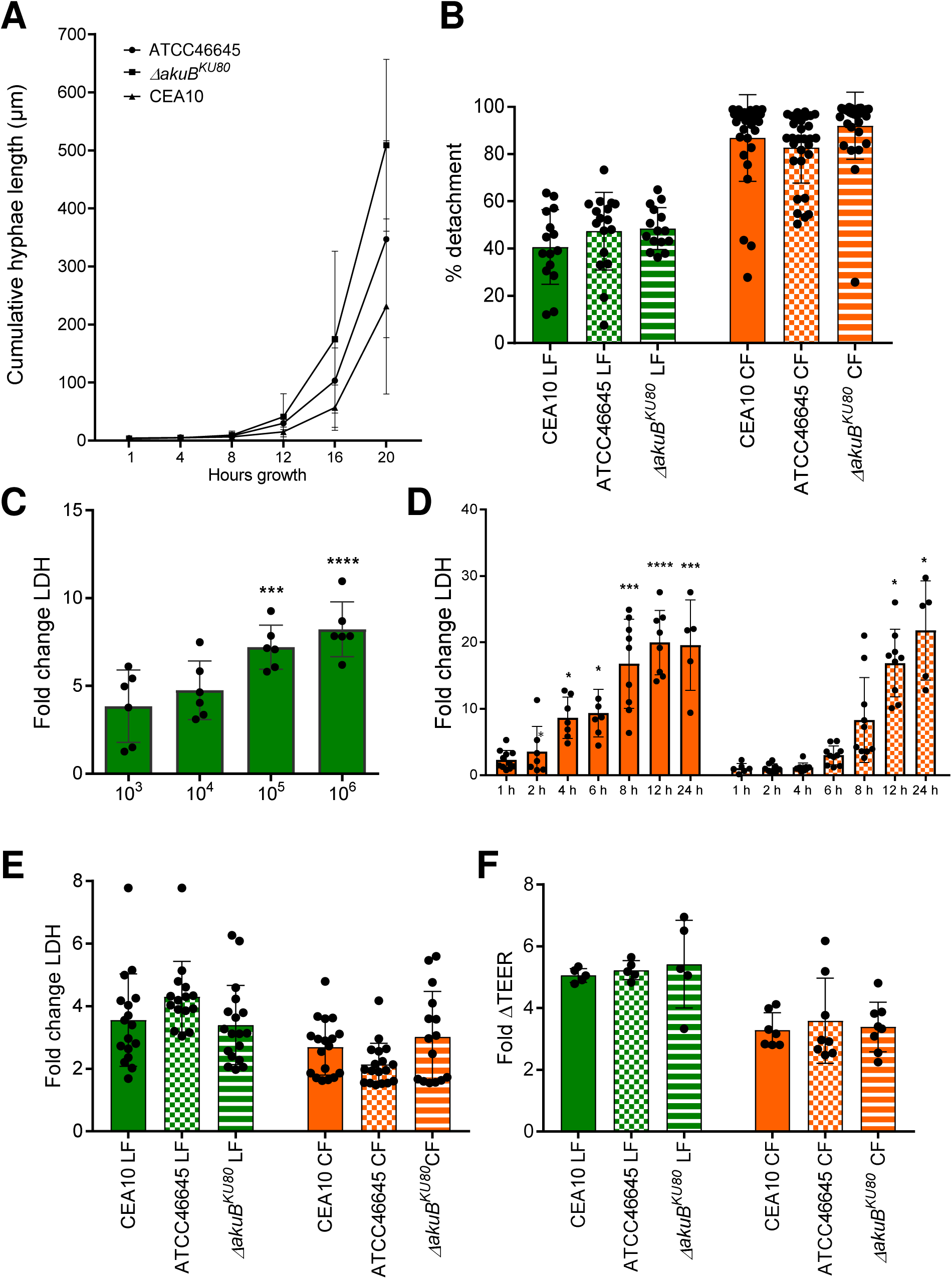
Temporal quantitative analysis of epithelial decay following challenge with live *A. fumigatus* spores or CF. **(A)** Cumulative hyphal growth (μm) of isolates CEA10, ATCC45546 and *Δaku*^*Ku80*^ after growth at 4, 8, 12, 16 and 20 h of incubation in supplemented RPMI. **(B)** Percentage of detachment of A549 cells following infection with *A. fumigatus* CEA10, ATCC45546 and *Δaku*^*Ku80*^ spores and CF^48^ thereof. **(C)** Fold change LDH release (relative to PBS challenge) following infection with the indicated doses of *A. fumigatus* CEA10 spores for 24 h. **(D)** Fold change LDH release (relative to PBS challenge) following challenge with a 5-fold or 10-fold diluted CF^48^ from CEA10 for indicated time points. **(E)** Fold change LDH release (relative to PBS challenge) of A549 cells following infection with *A. fumigatus* CEA10, ATCC45546 and *Δaku*^*Ku80*^ spores and CF^48^ thereof. **(F)** Fold change decrease in TEER between A549 monolayers incubated for 24 and 0 h with *A. fumigatus* CEA10, ATCC45546 and *Δaku*^*Ku80*^ spores and CF^48^ thereof. Data represent the mean of three biological replicates. Error bars show ± SEM. Data were analysed by non-parametric one-way ANOVA (Kruskal-wallis test) with Dunn’s multiple comparisons. Significance was calculated relative to challenge with vehicle control (PBS) unless otherwise stated. ^****^p ≤ 0.0001, ^***^p ≤ 0.001, ^**^p ≤ 0.01, and ^*^p ≤ 0.05

**Supplementary Figure 2:**
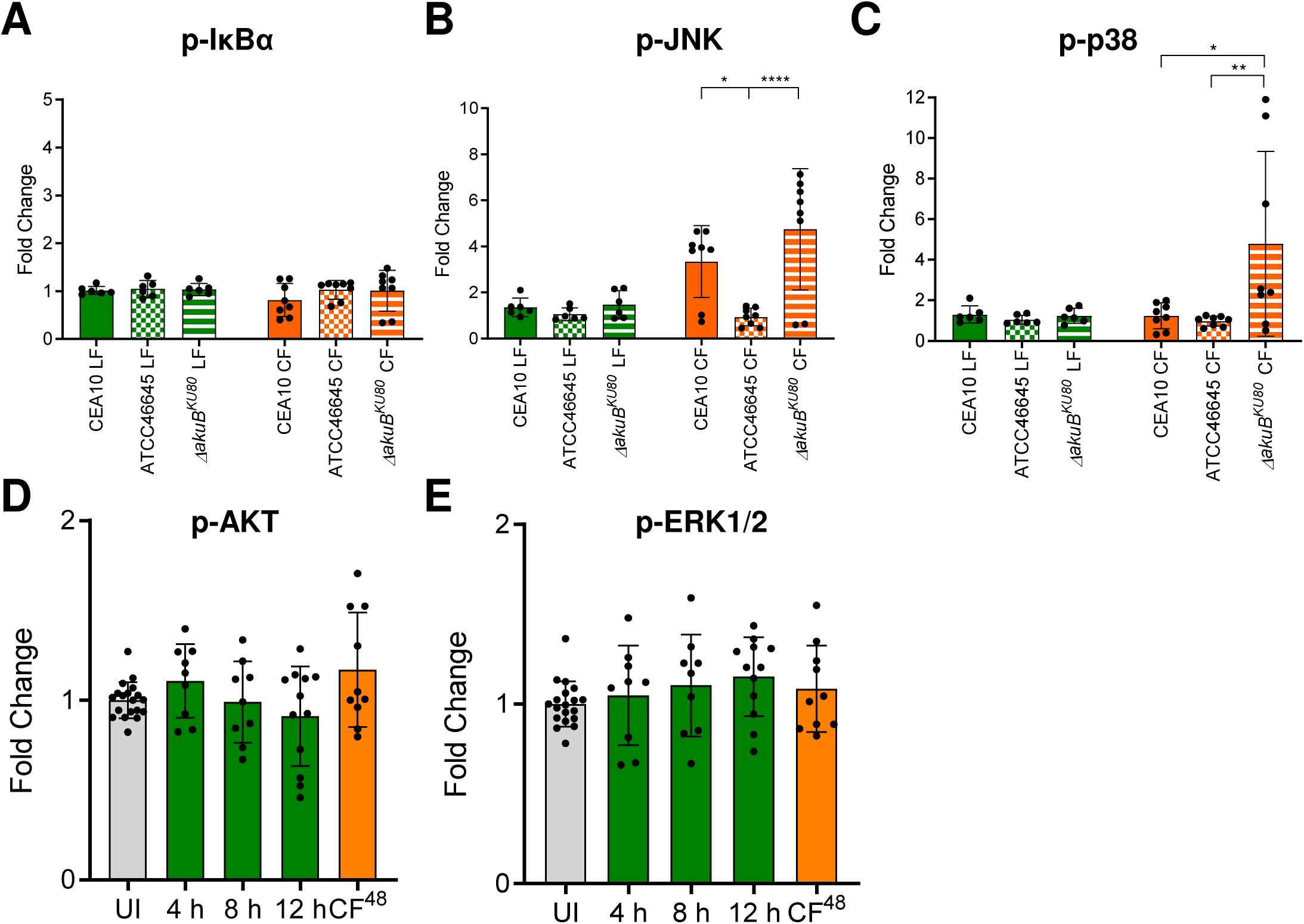
Different fungal morphotypes induce differential and dynamic phosphorylation and activation of host signalling proteins in A549 cells. **(A-C)** Fold change (relative to uninfected control, UI) phosphorylation of NF-kB (**A**: p-IkBα) and MAPK (**B**: p-JNK and **C**: p-p38) signalling following exposure to *A. fumigatus* CEA10, ATCC45546 and *Δaku*^*Ku80*^ spores (1×10^7^ spores/ml) for 12 h or respective 5-fold diluted CF^48^ for 4 h. **(D-E)** Fold change (relative to uninfected control, UI) phosphorylation of p-AKT **(D)** and p-ERK1/2 **(E)** following exposure to *A. fumigatus* spores (1×10^7^ spores/ml) for indicated time points or 5-fold diluted CF^48^ for 4 h. Data represent three biological replicates with 1-5 technical replicates. Error bars show ± SEM. Data was analysed by non-parametric one-way ANOVA (Kruskal-wallis test) with Dunn’s multiple comparisons. Significance was calculated relative to challenge with vehicle control (UI) and between each treatment as shown. ^****^p ≤ 0.0001, ^***^p ≤ 0.001, ^**^p ≤ 0.01, and ^*^p ≤ 0.05

**Supplementary Figure 3:**
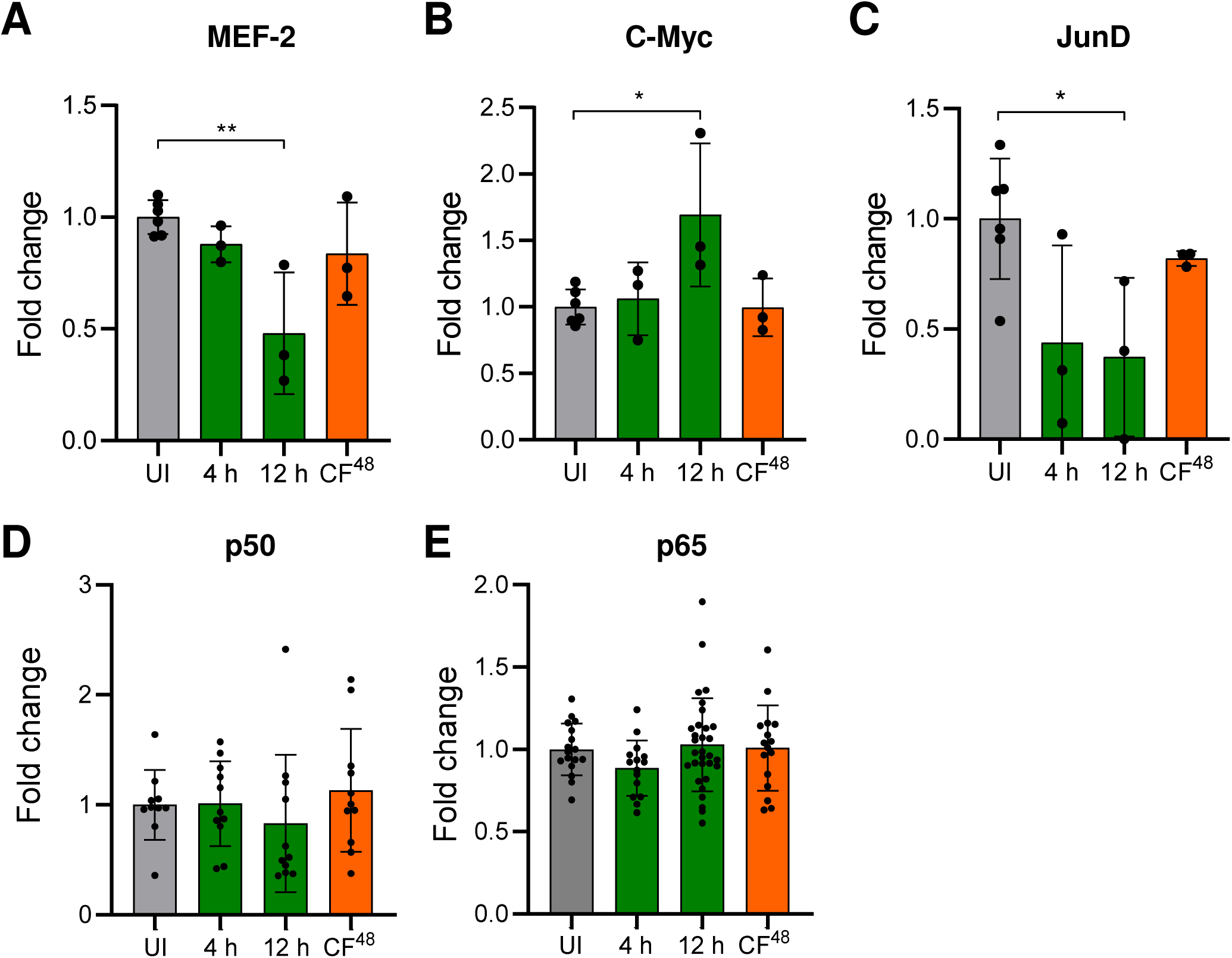
Different fungal morphotypes induce differential and dynamic activation of host transcription factors modulating epithelial host responses. **(A-E)** Fold change (relative to uninfected control, UI) in DNA binding activity of host transcription factors (**A**: MEF-2; **B**: c-Myc; **C**: JunD; **D**: p50 and **E**: p65) following exposure to *A. fumigatus* spores (10^7^ spores/ml) for indicated time points or 5-fold diluted CF^48^ (inoculum of 10^6^ spores/ml) for 4 h. Data represent three biological replicates with 1-5 technical replicates. Error bars show ± SEM. Data was analysed by non-parametric one-way ANOVA (Kruskal-wallis test) with Dunn’s multiple comparisons. Significance was calculated relative to challenge with vehicle control (PBS) as shown. ^****^p ≤ 0.0001, ^***^p ≤ 0.001, ^**^p ≤ 0.01, and ^*^p ≤ 0.05

**Supplementary Figure 4:**
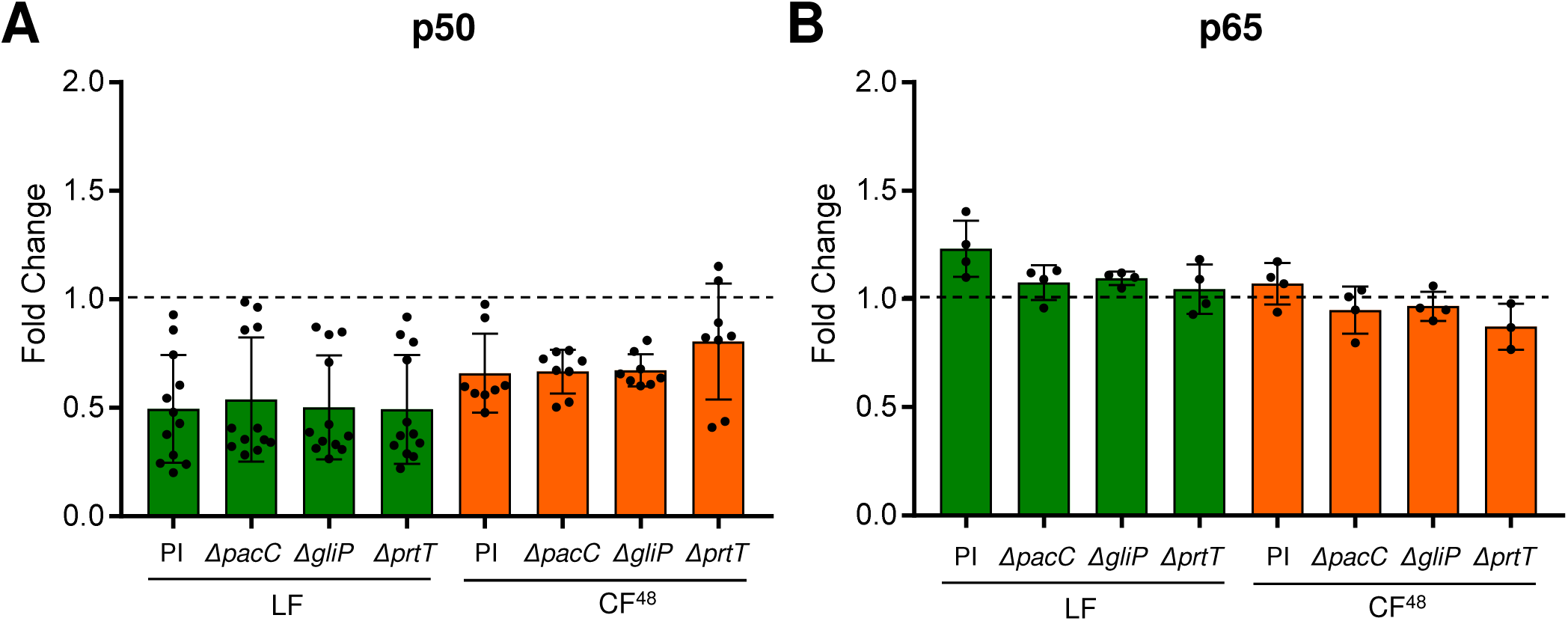
Deletion of Δ*pacC*, Δ*gliP* and Δ*prtT* does not impact p50 and p65 signalling upon challenge with *A. fumigatus* spores or CF. Fold change (relative to uninfected control, UI) in DNA binding activity of host transcription factors (**A**: p50 and **B**: p65) following exposure to *to A. fumigatus ΔpacC, ΔgliP and ΔprtT* mutants and respective parental isolate (PI) (10^7^ spores/ml) for 8 h or 5-fold diluted CF^48^ (inoculum of 10^6^ spores/ml) for 4 hours. Error bars show ± SEM. Data was analysed by non-parametric one-way ANOVA (Kruskal-wallis test) with Dunn’s multiple comparisons. Significance was calculated relative to challenge with vehicle control (PBS). ^****^p ≤ 0.0001, ^***^p ≤ 0.001, ^**^p ≤ 0.01, and ^*^p ≤ 0.05

## Supplementary Data 1

### *A. fumigatus* strains used in this study

| Isolate | Genotype | Reference/Source |
| --- | --- | --- |
| CEA10<br>(CBS 144-89) | Clinical isolate | (9) |
| $\DeltaakuB^{KU80}$ | $\DeltaakuBku80::pyrGAf-zeo$ | (46) |
| tdTomato<sup>ATCC46645</sup> | [ATCC46645];*his2A::Tdtomato* | (47) |
| $\Delta pacC^{A1160}$ | [A1160];$\DeltaakuBku80$; *pacC::Hyg* | (48) |
| $\Delta prtT^{A1160}$ | [A1160];$\DeltaakuBku80$; *prtT::Hyg* | (48) |
| $\Delta gliP^{CEA17}$ | [CEA17];$\DeltaakuBku80::pyrGAf$; *gliP::(neo-A.n.pyrG-neo)* | (27) |

## Supplementary Data 2

### DAPI Counter macro to automatically process and count DAPI objects

// DAPI Counter macro to automatically process and count DAPI objects

// This macro is optimised for wide-field imaging with a 20x 0.75NA objective lens and a 6.45um pixel size camera, giving a digital pixel size of 0.323um. If your images are of a different resolution please contact for assistance

// This macro was written by Darren Thomson June 2016. Any comments or quereys should be directed to

// Instructions: Create a folder with only the fluorescent images of DAPI + create a folder to deposit the processed outlines images

// Drag and drop this macro file onto FIJI and ‘Run’ the script. OR create a new macro and paste this code into the window and ‘Run’ the script.

// Direct the macro to the input folder with all the DAPI images are + the output folder to deposit processed images

// Save the results data to Excel from the macro and arrange/illustrate the data accordingly in Excel.

dir1 = getDirectory(“Where are the DAPI images”);

format = getFormat();

dir2 = getDirectory(“Where will I save the processed images?”);

list = getFileList(dir1);

setBatchMode(true);

for (i=0; i<list.length; i++) {

showProgress(i+1, list.length);

open(dir1+list[i]);

run(“Subtract Background…”, “rolling=35”);

run(“Gaussian Blur…”, “sigma=3”);

setAutoThreshold(“Huang dark”);

//run(“Threshold…”);

run(“Convert to Mask”);

run(“Watershed”);

run(“Analyze Particles…”, “size=400-5000 circularity=0.50-1.00 show=Outlines summarize”);

if (format==“8-bit TIFF” || format==“GIF”)

convertTo8Bit();

saveAs(format, dir2+list[i]);

close();

}

function getFormat() {

formats = newArray(“TIFF”, “8-bit TIFF”, “JPEG”, “GIF”, “PNG”,

“PGM”, “BMP”, “FITS”, “Text Image”, “ZIP”, “Raw”);

Dialog.create(“Batch Convert”);

Dialog.addChoice(“Convert to: “, formats, “TIFF”);

Dialog.show();

return Dialog.getChoice();

}

function convertTo8Bit() {

if (bitDepth==24)

run(“8-bit Color”, “number=256”);

else

run(“8-bit”);

}

## Supplementary Data 3

### RIPA Buffer recipe

(0.05 M (50 mM) Tris-HCl pH7.5, 0.15M (150 mM) NaCl, 1% Triton X-100, 1% Sodium Deoxycholate, 0.1% SDS and 20 mM EDTA) with 10 μl/ml protease (Cat. No 78410, Thermofisher, United Kingdom) and 10 μl/ml phosphatase (Cat. No P5726-5ML, Sigma, United Kingdom) inhibitors.

## Supplementary Data 4

Cytokine profile array in response to live *A. fumigatus* and CF Submitted as Excel spreadsheet

### Human XL cytokine (HXL) profiling procedure

To determine the global cytokine expression from epithelial cells in response to infections with live conidia and CF, cell free culture supernatant was analysed using the HXL cytokine profile kit from R&D systems according to the manufacturer’s instructions. Briefly, a nitrocellulose membrane, pre-coated with same amount of capture antibody against the cytokines of interest was incubated with a blocking solution on a rocking platform for 1 h. Culture supernatant from each treatment was diluted 1:3 to a final volume of 1.5 ml. The membrane was incubated with the culture supernatant overnight at 2-8°C on a shaker. Membrane was washed 3x on a shaker for 10 min each and then incubated with detection antibody cocktail in specified diluents for 1 h. The membrane was washed as before and incubated with a 1:2000 dilution of streptavidin-phycoerythrin conjugate for 30 min. The membrane was washed and developed quickly using a 1:1 ratio of chemi-reagent 1 and 2 on a plastic sheet for 1 min at room temperature, protected from light. Excess reagent mix was removed and the membrane was covered with a plastic wrap and exposed to X-ray film for 1-10 min using a ChemoDoc MP imaging system (BIO-RAD). The relative amounts of cytokines were quantified by mean pixel density of the blots using ImageJ software and micro-array profile plugins (R&D) and normalized to the positive and negative control blots first and then to the PBS treated controls.

